# The relationship between benthic nutrient fluxes and bacterial community in Aquaculture Tail-water Treatment Systems

**DOI:** 10.1101/2021.08.18.456883

**Authors:** Regan Nicholaus, Betina Lukwambe, Wen Yanga, Zhongming Zhenga

## Abstract

Constructed-wetlands, Biofilms, and sedimentation are potential aquaculture tail-water treatments however their roles on the distribution of benthic microbial community and the way they affect the interaction between microbial community and inorganic nutrient fluxes have not been fully explored. This study applied 16S rRNA high-throughput sequencing technology to investigate the microbial community distribution and their link with nutrient fluxes in an aquaculture tail-water bioremediation system. Results showed that bacterial community compositions were significantly different in constructed-wetland and biofilm treatments (p<0.05) relative to sedimentation. The composition of the 16S rRNA genes among all the treatments was enriched with *Proteobacteria, Bacteroidetes, Firmicutes*, and *Flavobacteria*. NMDS analysis showed that the bacterial composition in constructed-wetland and biofilm samples clustered separately compared to those in sedimentation. The Functional-Annotation-of-Prokaryotic-Taxa analysis indicated that the proportions of sediment-microbial-functional groups (aerobic-chemoheterophy, chemoheterotrophy, and nitrate-ammonification combined) in the constructed-wetland treatment were 47%, 32% in biofilm and 13% in sedimentation system. Benthic-nutrient fluxes for phosphate, ammonium, nitrite, nitrate and sediment oxygen consumption differed markedly among the treatments (p<0.05). Canonical correspondence analysis indicated constructed-wetland had the strongest association between biogeochemical contents and the bacterial community relative to other treatments. This study suggests that the microbial community distributions and their interactions nutrient fluxes were most improved in the constructed-wetland followed by the area under biofilm and sedimentation treatment.

## Introduction

Intensive aquaculture farming practices have contributed overwhelming allochthonous organic matter (OM), excreta, food-wastes, and dissolved nutrients (e.g., nitrogen and phosphorous) with substantial impacts on the environment [1–6]. The pollution increment within the environment has increased the concern for the adoption of aquaculture effluent treatments. Biological effluent treatments such as constructed wetland and biofilm and physical treatments like sedimentation are highly used in treating aquaculture effluents [7–9].

Constructed-wetlands are artificially designed biological systems consisting of a complex substrate, plants (macrophytes), microbes, and water bodies forming an ecosystem [10]. Besides they are a well-established, viable, suitable, and cost-effective method for treating various forms of wastewater such as industrial and agricultural wastewaters [11]. Wetland plants photosynthesize by their aboveground organs, while their roots and rhizospheres interact with the below-sediment to drive the productivity of the heterotrophic soil biota [12, 13]. Wetland rhizospheres are essentially oxic-habitats or niches created by the roots’ aeration and can markedly affect the diversity of the wetland’s heterotrophic biota [14]. Wetlands can influence water and/ or sediment physicochemical properties through different mechanisms including microbial OM mineralization, sedimentation and substrate-adsorption processes [9,15].

Biofilms are ubiquitous and auto-aggregate forms of heterogeneous microbial communities that are attached to each other and can invariably develop on solid surfaces exposed to aquatic environments [16, 17]. Biofilm communities consisted of bacteria and microalgae which secrete an extracellular polymeric substance matrix (polysaccharides) which facilitates the adhesion of the community to other substrates. The physical nature of biofilm exopolymers has a great adsorptive capacity with a super binding affinity for nutrients [17]. Biofilms can contribute substantially to nutrient cycling, organic matter degradation, and community enrichment due to bacteria mineralization [9,18, 19].

Sedimentation is the physical process by which suspended material such as clay, silts and other organic particles found in the water settle by gravity. The resulting sedimentary niche at the settling area could form microbial communities and nutrient-rich ecosystems [20]. Provoost et al. [21] reported that sediment can harbor up to 30% of the pelagically produced organic matter. Various pollutants in the water body are deposited onto the sediments and through microbial waste degradation processes such as bioremediation undergo biological transformations resulting in increased nutrient cycling, pollution reduction, and bacterial diversity [22–24]. Sediment microorganisms like heterotrophic marine bacteria are very crucial in nutrients cycling and OM processing [25].

The strength and efficacy of the constructed wetland and biofilm treatments are supported by the consortium of bacterial communities [9, 26]. Thus, exploring the relationship between biological treatment methods such as constructed wetlands and biofilms on bacterial community and sediment properties is imperative. A couple of studies investigated various effluent treatment systems focusing on microbial community composition and distribution [9, 19, 27]. However, the impacts associated with constructed wetland, biofilm and sedimentation in response to benthic properties, nutrient fluxes, and distribution of bacterial community during bioremediation of aquaculture tail-water remain unclear. In this study, we aimed to (i) evaluate the distribution of microorganisms in constructed wetland, biofilm, and sedimentation (ii) explore the distribution of microbial functional groups among the treatments and (iii) investigate the relationship between nutrient fluxes and microbial functional groups/microbial community. This study will add knowledge on the distributions of sediment microbial community and nutrient fluxes of an aquaculture tail-water treatment system

## Materials and Methods

### Study area, Experimental design and Sampling

The experiment was conducted in Ningbo Xiangshan Bay, Zhejiang Province, China, at a land-based aquaculture tail-water treatment system constructed. This system was primarily to restore effluents resulting from an intensive Commercial Vannamei Shrimp (*Litopenaeus vannamei*) production farm. A comprehensive aquaculture tail-water treatment system composed of subsystems: constructed-wetland, biofilm, and sedimentation was studied [28]. The constructed-wetland subsystem was composed of emergent macrophytes, *Spartina anglica*, occupying 400 sq. m of the total system area. The planting densities of *S. anglica* were 50% of the total wetland cover. These plants grew rapidly to colonize the wetland and they were not harvested during this study. The biofilm system was deployed with suitable aeration facilities and suspended carriers in the form of fiber threads (adhesive matrix of extracellular polymeric substances) for enhancing the surface area for microorganism attachment. The physical sedimentation consisted of bare sediment surface, overlying aquaculture effluent water, and aeration. This system was involved in filtering and settling large particles of the incoming effluent water from the production center.

Sampling started one year after the project launched to let the ecological succession develop. To ensure a representative sampling strategy, data was collected three times consecutively, between April to July. Four different sampling points from each system were identified and sampled. 0.5L of the overlying water was collected from each system for water quality analysis. Using a handheld sediment corer four undisturbed sediment cores (8 cm height) from each system were gently collected in cylindrical plastic tubes (i.d. 6.4 cm, height 19.4 cm). The sediment cores and water samples were immediately brought back to the laboratory for physicochemical analysis and incubation. Water samples were stored at 4°C, whereas the sediment cores were kept ready for the incubation experiment.

The incubation experiment was done as previously described [29]. Water samples for the determination of benthic flux rates for the total ammonia nitrogen (TAN), nitrate (NO_3_^−^-N) and nitrite (NO_2_^−^-N), and soluble reactive phosphate (SRP) were collected, filtered in 0.45 GF/F and stored under −20°C until analysis. After the incubation experiment, using a clean stainless steel microspatula, the sediment cores were sliced into three sub-sampling points (surface 0-2 cm, middle 2-4 cm, and bottom 4-8 cm). These subsamples were thoroughly homogenized and divided into two portions. One potion was freeze-dried for physicochemical contents analysis and the other potion was stored in clean polypropylene tubes at −20°C for the 16S rRNA extraction.

### Analysis of physicochemical contents and nutrient flux rates

The physicochemical water parameters (dissolved oxygen, temperature, and salinity) were measured *in situ* during sampling using a handheld automated YSI 6000 multi-parameter probe (USA). All water samples were analyzed using standard methods [30], where TAN was treated with indophenol blue, NO_2_^−^-N/NO_3_^−^-N with the cadmium-copper reduction and the SRP were treated with the ammonium molybdate/ascorbic acid method. All the concentrations of inorganic nutrients were measured using a WESTCO SmartChem discrete analyzer 200 USA. Nutrient flux rates (μmol m^−2^h^−1^) and SOC were calculated from slopes of a linear regression concentration against time using the equation previously described [29].

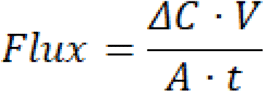

Where: Flux is the nutrients or sediment oxygen fluxes (mmol m^−2^h^−1^); ΔC (mgL^−1^) is the change in concentration of oxygen/nutrients (prior and after incubation); V (m^3^) the volume of overlying water; A (m^2^), is the cross-sectional area of the incubation chamber; t (h) is the duration of incubation.

The sediment grain size distribution was determined using sieves with different mesh sizes [31]. Briefly, grain-size parameters were conducted mechanically from oven-dried subsamples using standard sieving methods for the sand content (500-63 μm) and sedigraph techniques for the silt/clay fraction (<63 μm). Particles sizes (clay: <0.002, silt: <0.02, fine sand: <0.2, sand: <2) were determined. Sediment OM was determined using the loss on ignition method (LOI) [32]. The sediment samples were freeze-dried, pulverized, and pre-weighed before being placed in a muffle furnace at 475°C for 4 h. Then the samples were reweighed with the difference equals to the %OM content. TOC (total organic carbon), TON (total organic nitrogen), and C/N (carbon/nitrogen) ratio were analyzed commercially by using Carbon Elemental Analyzer. Briefly, during pretreatment, 5g of the post-freeze-dried wet-sediment were ground using a mortar into powder to pass through a 1-mm mesh sieve. Before analysis, further pretreatment procedures necessary especially for TOC were done to remove the carbon dioxide by adding 1:1 HCl and oven-dried at 80°C, overnight to a constant weight.

### Extraction, amplification and MiSeq sequencing of the benthic bacterial DNA

The total genomic sediment DNA extraction was performed from ~0.5g of homogenized sediment samples using the PowerSoil™ DNA isolation kit (MoBio Laboratories, Inc., USA) according to the manufacturer’s recommendations. The extracted genomic DNA was stored at −80°C until amplification. The PCR amplification conditions were according to Lukwambe *et al*. [28] whereby the total genomic DNA was dissolved in 100 μl of DES, supplied with the kit. The DNA quality and/or quantity of the samples were measured using a spectrophotometer (NanoDrop Technologies Inc., Wilmington, DE, USA) at the A260/A280 ratio. A combination of reverse primer (5′-GGACTACHVGGGTWTCTAAT-3′) and a forward primer (5′-CCTACGGGAGGCAGCAG-3′) for the hypervariable V3-V4 regions of the 16S rRNA gene was used. 5 μl of the total DNA template was used and amplified in a 50-μl reaction system. Then the amplification process followed 30 cycles of 95°C denaturation for 30 s, annealing (55°C, 30 s), and extension (72°C, 45 s) and a final extension for 5min at 72°C. Successful amplification product and size of the PCR was electrophoresed in 1% agarose gel. The triplicate amplified products of each sample were pooled, purified, equilibrated, and sequenced in an Illumina MiSeq high-throughput sequencing platform.

### Bioinformatics analysis

The sequencing process of the paired reads was initially joined with FLASH using default settings [33], then, the Raw FASTQ files were processed using Quantitative-Insights-Into-Microbial-Ecology (QIIME version 1.8.0, [34]. The operational taxonomic units (OTUs) assigned at a 97% similarity cut-off point in all samples were clustered using USEARCH (version 7.1, http://drive5.com/uparse/). The sequences were quality filtered based on sequence length, quality score, chimera, and primer mismatch thresholds. In a nutshell, homopolymer runs exceeding 6 bp were screened-out by PyroNoise. Sequences with the same barcodes were assigned to the same sample. The phylotypes were performed using the UCLUST algorithm [35]. The most abundant sequences of each phytotype were selected as the clean sequence and were taxonomically assigned (Greengenes database, release 13.8) using PyNAST [36]. Diversity indices (Shannon index, Simpson, Chao1, and observed OTUs) were performed using the phylogenetic tree (QIIME pipeline).

### Statistical analyses

The variations of the different physicochemical variables were analyzed by a one-way or two-way repeated ANOVA. Post Hoc tests were performed to determine the significant groups. The normal distribution and homogeneity of variances among treatments were verified before the ANOVA test. All the data were Hellinger transformed post statistical analyses, and then normalized by using the function decostand/p-p plot in the “vegan” package to improve normality and homoscedasticity. Permutational multivariate analysis of variance-PERMANOVA (Bray-Curtis dissimilarity matrices) [37], phyloseq v1.22.3 and Nonmetric Multidimensional Scaling (NMDS) was performed to analyze the microbial community composition among the treatments. One-way analysis of similarity (ANOSIM) was used to verify whether the distribution of different samples visualized in the NMDS plot was significant [38]. Canonical correspondence analysis (CCA) was used to analyze the correlations between bacterial community compositions and environmental variables.

The sediment microbial functional groups were predicted by using the Functional Annotation of Prokaryotic Taxa (FAPROTAX) database. According to Louca *et al*. [39], the annotated bacterial OTU table from the Silva database was read, and the data was matched with the species information in the database using a python program. The predicted functions were outputted through FAPROTAX (http://www.ehbio.com/ImageGP/). The annotation results were used to describe the microbial functional compositions and abundance of related metabolic pathways involved in ammonification, denitrification, carbohydrate metabolism, aromatics degradation, and nitrogen fixation. The relative abundances of the functional groups were calculated as the cumulative abundance of OTUs assigned to each functional group, which was obtained by standardizing the cumulative abundance of OTUs correlated with at least one function. All statistical analyses were performed with R, (version, 3.6.1) [40] and the results of the statistical tests were considered to be significant at p≤ 0.05. The figures were drawn with R and OriginPro 8.0 software. **Data deposition:**The sequences used in this study have been deposited in the GeneBank of NCBI with the BioProject database ID PRJNA593691 (https://www.ncbi.nlm.nih.gov/sra/PRJNA593691) and SRA accession numbers ranging from SAMN13483434 to SAMN13483469.

## Results

### Sediment and water physicochemical contents

The sediment physicochemical contents are described in Table 1. The results indicate distinct differences in sediment organic and inorganic contents among the systems. The surface sediments of the constructed-wetland consisted of 79% medium sand, 17% very fine sand, and 4% silt/clay whereases the compositional contents in the biofilm were 68% medium sand, 25% very fine sand, and 7% silt. The sedimentation system was dominated by 84% (medium sand), 11% (very fine sand), and 5% (silt). Cores from the sedimentation system had significantly higher contents of OM, TN, TP, TOC, and C/N ratio at all three depths (0-2 cm, 2-4 cm, and 4-8 cm), relative to the others. C/N ratio were 8.17±1.5 (biofilm), 7.7±1.6, (constructed-wetland) and 12.32±3.1 (sedimentation) on average. Total OM was much lower in biofilm (depth 0-2 cm) compared to constructed-wetland and sedimentation. All sediment organic contents varied differently between the systems however no stable variational trends were observed within different depths of the same treatment (Table 1, p>0.05).

**Table 1.** Mean (±SD) values of the sediment organic contents among the treatments (Constructed-wetland, Biofilms, and Sedimentation). OM represents organic matter, TN: Total nitrogen, TP: Total phosphorus, TOC: Total organic carbon and C/N: Carbon to nitrogen ratio

| System | Depth (cm) | OM% | TN% | TP% | TOC% | C/N |
| --- | --- | --- | --- | --- | --- | --- |
| **CW** | Surface | $3.13 \pm 0.14$ | $0.48 \pm 0.05$ | $0.031 \pm 0.061$ | $2.94 \pm 0.06$ | $6.12 \pm 1.31$ |
| | Middle | $3.23 \pm 0.11$ | $0.38 \pm 0.02$ | $0.029 \pm 0.053$ | $3.13 \pm 0.03$ | $8.24 \pm 1.05$ |
| | Bottom | $3.07 \pm 0.08$ | $0.30 \pm 0.12$ | $0.021 \pm 0.006$ | $2.63 \pm 0.04$ | $8.77 \pm 2.09$ |
| **BF** | Surface | $1.57 \pm 0.05$ | $0.43 \pm 0.03$ | $0.086 \pm 0.04$ | $3.10 \pm 0.06$ | $7.21 \pm 0.81$ |
| | Middle | $2.55 \pm 0.04$ | $0.41 \pm 0.01$ | $0.081 \pm 0.03$ | $3.07 \pm 0.02$ | $7.49 \pm 3.32$ |
| | Bottom | $3.47 \pm 1.02$ | $0.22 \pm 0.01$ | $0.952 \pm 0.02$ | $2.16 \pm 0.08$ | $9.82 \pm 3.74$ |
| **SD** | Surface | $4.79 \pm 0.26$ | $0.16 \pm 0.01$ | $0.263 \pm 0.01$ | $2.31 \pm 0.06$ | $14.43 \pm 5.64$ |
| | Middle | $4.12 \pm 0.24$ | $0.21 \pm 0.01$ | $0.289 \pm 0.041$ | $2.33 \pm 0.25$ | $11.09 \pm 4.81$ |
| | Bottom | $3.98 \pm 0.31$ | $0.26 \pm 0.12$ | $0.204 \pm 0.025$ | $2.98 \pm 0.22$ | $11.46 \pm 6.06$ |

### Nutrients flux rates among the treatments

All dissolved inorganic nutrient flux rates showed an efflux trend among the treatments. The mean release rates of TAN, NO_3_^−^-N, NO_2_^−^-N, and SRP fluxes between biofilm, constructed-wetland, and sedimentation cores were significantly different (2-way ANOVA, p<0.05). NO_2_^−^-N and TAN accounted for more than 87.51% (constructed-wetland) and 71.14% (biofilm) net flux rate relative to 37.43% (sedimentation) (Fig 1). The NO_3_^−^-N flux rates were 396.15±61.09 μmol m^−2^h^−1^ (constructed wetland), 249.83±71.12 μmol m^−2^h^−1^ (biofilm), 173.7±33.01 μmol m^−2^h^−1^, (sedimentation) (Fig 1B). The constructed wetland had the highest exchange rate of NO_3_^−^-N, and NO_2_^−^-N relative to other systems (biofilm and sedimentation). The SRP had the highest mean flux rate in biofilm. The release rate of TAN into the overlying water (constructed-wetland) was approximately twice higher in both biofilm and sedimentation, indicating sedimentary remineralization of ammonia and nitrate. SOC were 4.91±0.75 mmol m^−2^h^−1^ (biofilm), 3.82±0.37 mmol m^−2^h^−1^ (constructed wetland), 1.89±0.31 mmol m^−2^h^−1^ (sedimentation) (Fig 1A). Oxygen level in sedimentation subsystem was the lowest followed by constructed wetland and finally biofilm. Generally, the mean release rates of all nutrient groups including soc followed the order: biofilm>constructed wetland>sedimentation.

**Fig 1.**
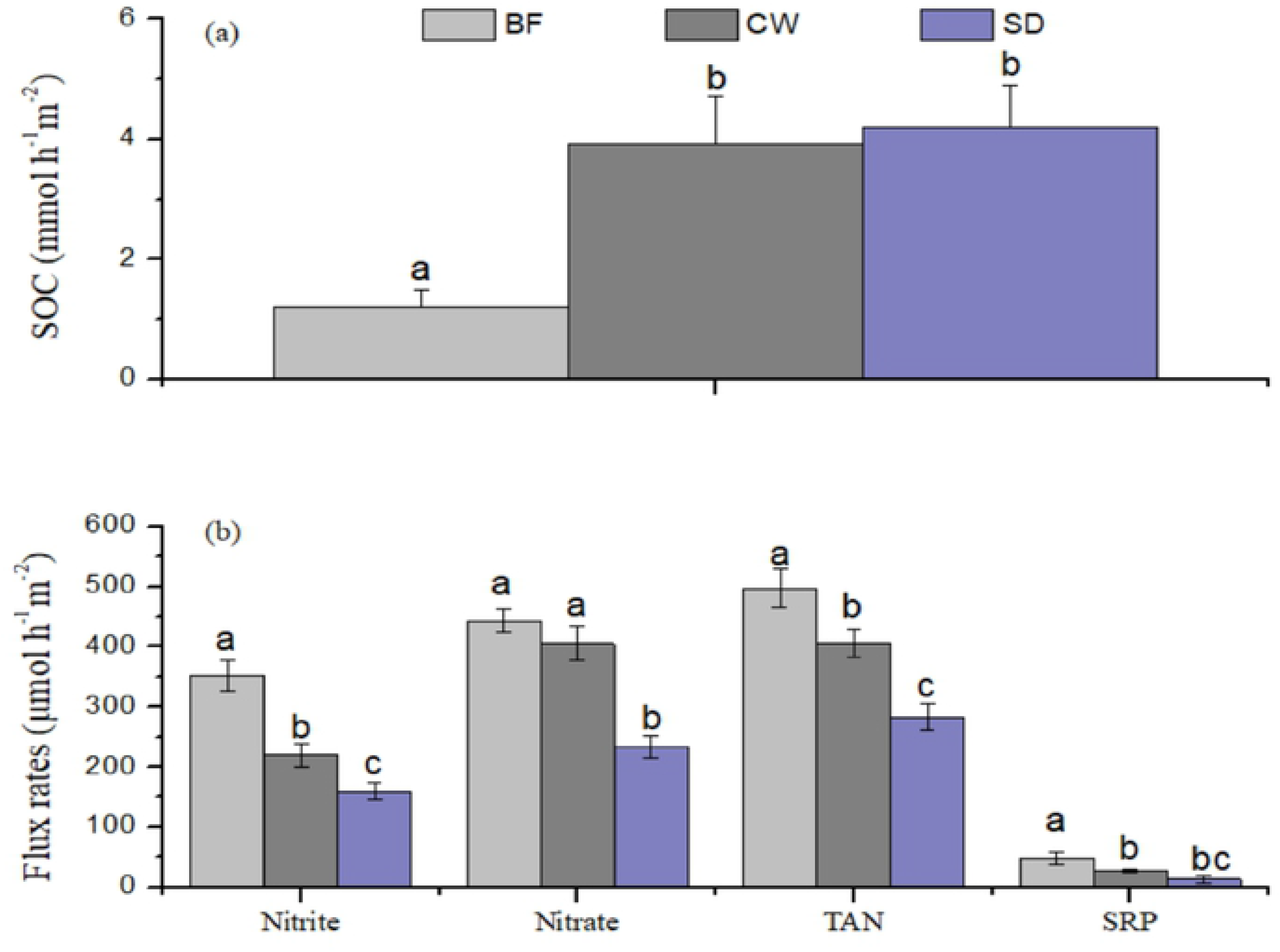
Benthic inorganic nutrient flux rates. (A) Sediment oxygen consumption-SOC and (B) (Total ammonia nitrogen-TAN, nitrate, nitrite and soluble reactive phosphate-SRP) estimated during laboratory incubations (mean±SD, n=3) in each treatment system

### Microbial community composition and structure

The bacterial community compositions varied among depths and between the treatment systems (Fig 2). A total of 519, 692, and 837 OTUs were identified for sedimentation, constructed-wetland, and biofilm treatments respectively. Jointly the OTUs represent 54 phyla, 85 classes, 152 families, and 471 genera among all treatments. The relative content of the microbial community (>0.3% relative abundance) at phylum, class, and family level is illustrated (Fig 2A-C). At the phylum level, *Proteobacteria* were the most dominant community in all three systems accounting for 32.17±7.51%, followed by *Bacteroidetes* (29.32±7.04%), *Chloroflexi* (20.65±6.21%), *Actinobacteria* (19.44±5.92%), *Firmicutes* (13.92±4.09%), *Acinetobacter* (11.85±3.71%) and *Planctomycetes* (8.83±2.54%) (Fig. 2A). The phylum *Firmicutes* and *Proteobacteria* were most dominant in constructed wetland and biofilm, while the sedimentation community was mainly dispersed by *Firmicutes, Proteobacteria*, and *Bacteroidetes*. *Gamma-, Delta-*, and *Alpha-proteobacteria* were dominant classes in all systems, followed by *Anaerolineae*, *Actinobacteria*, *Cytophagia*, *and Flavobacteriia* (Fig 2B). Further, at the family level, several predominantly expressed bacterial taxa ((*Clostridiaceae* and *Acidaminobacteraceae*, (Order*-Clostridiales*)*, Rhodobacteraceae* (Order*-Rhodobacterales*) *Chloroflexi, Anaerolineaceae*, [*Thermodesulfovibrionaceae*] (order*-Nitrospirales*) were predominant in all three treatments (Fig 2C). The family *Flavobacteriaceae* was highly distributed in biofilm (15 to 65-fold) relative to constructed-wetland (9-31-fold) and sedimentation (7-17-fold). Other families were *Nitrospiraceae* and *Planctomycetaceae* with a 20-50-fold higher (biofilm) relative to constructed-wetland and sedimentation (jointly 6-12-fold). The distribution of the most dominant bacterial community (at the genera level) among the treatments is represented by heatmap (Fig 3). The heatmap includes the top thirty genera, which represent 94.2-97.5% of all 16S rRNA bacterial genes reads. The *Disulvococcus, Novosphingomium, Fusibacteria, Kordia, Clostridium*, and *Lysobacter* were the genera highly distributed among the treatments (Fig 3; 0-4 cm depth). Vertically, the proportions of *Proteobacteria, Acidobacteria*, and *Bacteroidetes* were high in surface sediments, whereas *Chloroflexi* and *Firmicutes* tended to be enriched in deep layers.

**Fig 2.**
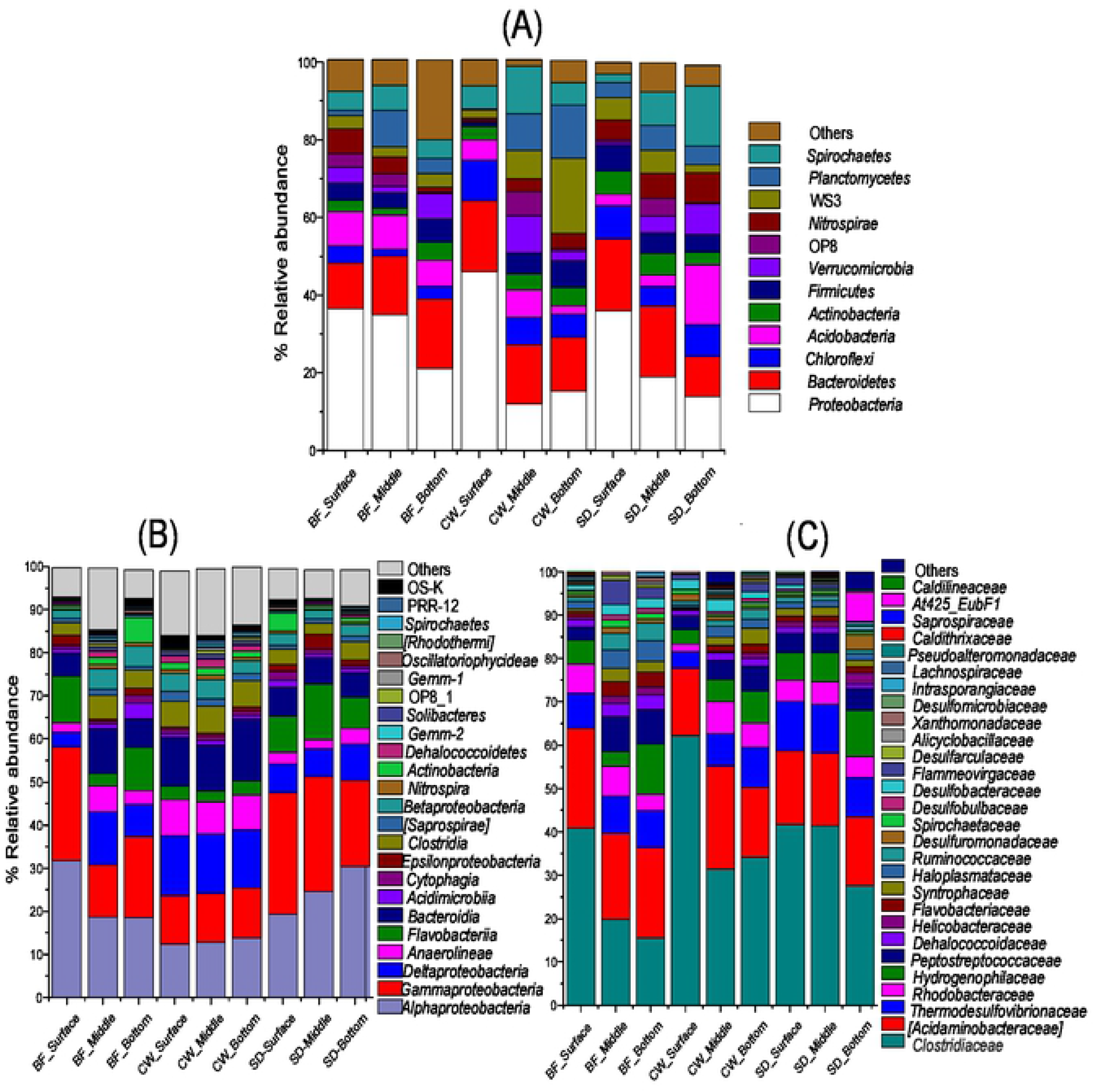
The relative abundance of the vertical sediment microbial community composition and diversity among the treatment systems (Biofilm, Constructed-wetland, Sedimentation) at the phylum (A), class (B) and family (C) levels revealed by 16S rRNA genes sequencing. Taxa making up less than 0.03% of total composition in all libraries were classified as ‘others’

**Fig 3.**
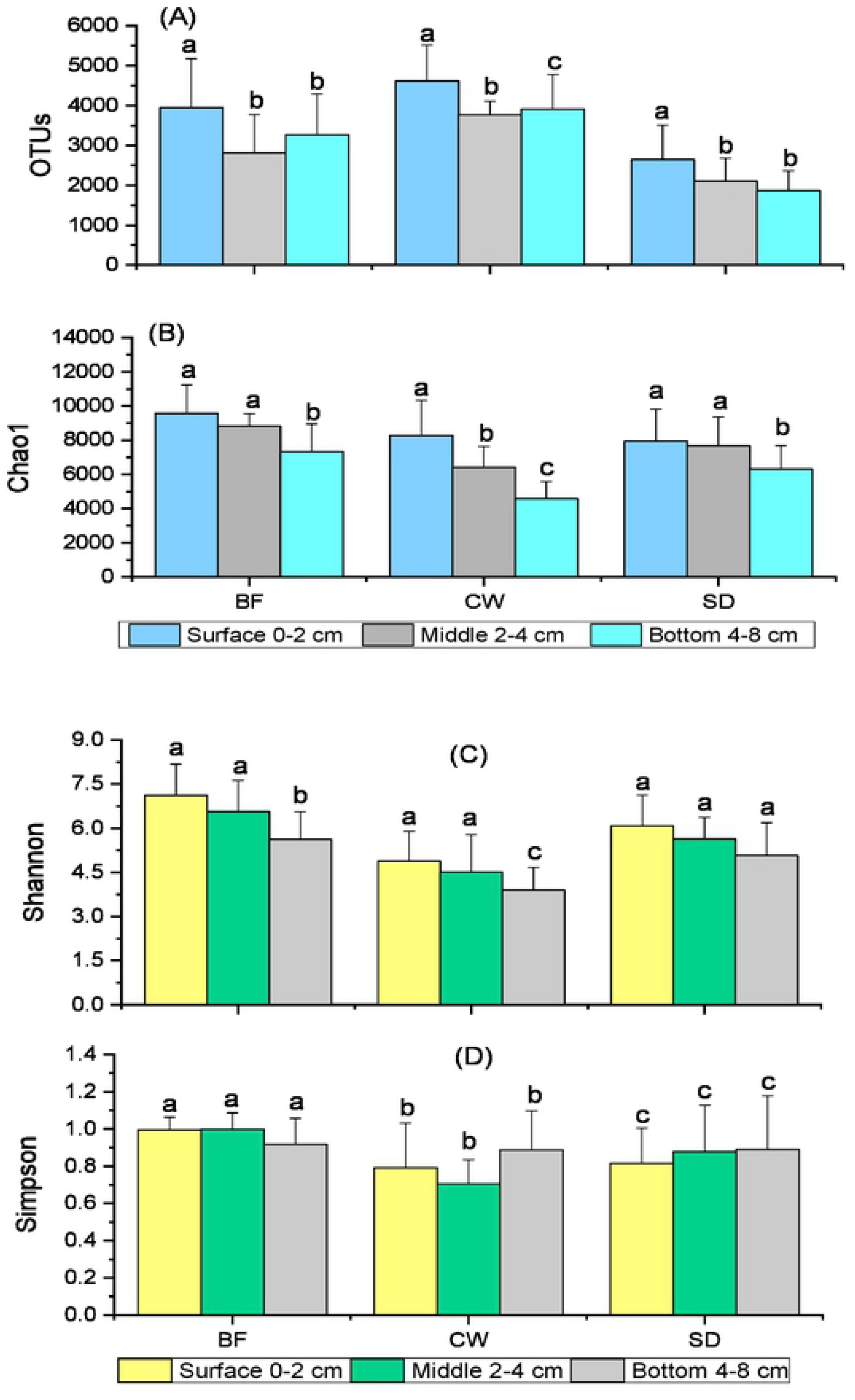
Alpha diversity estimates of each treatment at different sampling depths obtained by 16S rRNA genes high-throughput sequencing. (A) Number of OTUs, (B) Chao1 richness estimate index, (C) Shannon index and (D) Simpson

### Diversity of bacterial community among the systems

The bacterial community differed significantly among the treatments (ANOSIM, p = 0.031). The microbial community richness estimate (Chao1) ranged from 7321 to 9531 sequences (biofilm), 4637 to 9017 (constructed-wetland), 6214 to 8973 (sedimentation). Biofilm had significantly Chao1 values (p<0.05) and constructed-wetland had the highest bacterial diversity (Shannon index values). Bacterial richness estimates were highest and most diverse at the surface sediments (0-2 cm) and dropped with a depth increase (Fig 3). The Shannon index ranged from 6.79 (surface), 6.03 (middle) to 5.81 (bottom) in biofilm samples and 6.21 (surface), 6.05 (middle), 4.93 (bottom) in constructed-wetland samples whereas sedimentation ranged from 5.06 (surface), 3.5 (middle), 2.53 (bottom). The diversity trend order was constructed-wetland>biofilm>sedimentation. Moreover, the NMDS ordinations assessment revealed marked differences in community composition grouping patterns between the systems, and between depths (Fig 4). The samples were grouped separately within depths and between treatments. The microbial community in constructed-wetland treatment samples was more clearly separated suggesting the highest species dissimilarity compared to other treatments. PERMANOVA further confirms that bacterial community composition between the treatments and within depth groups was significantly different (Table 2: p<0.05).

**Fig 4.**
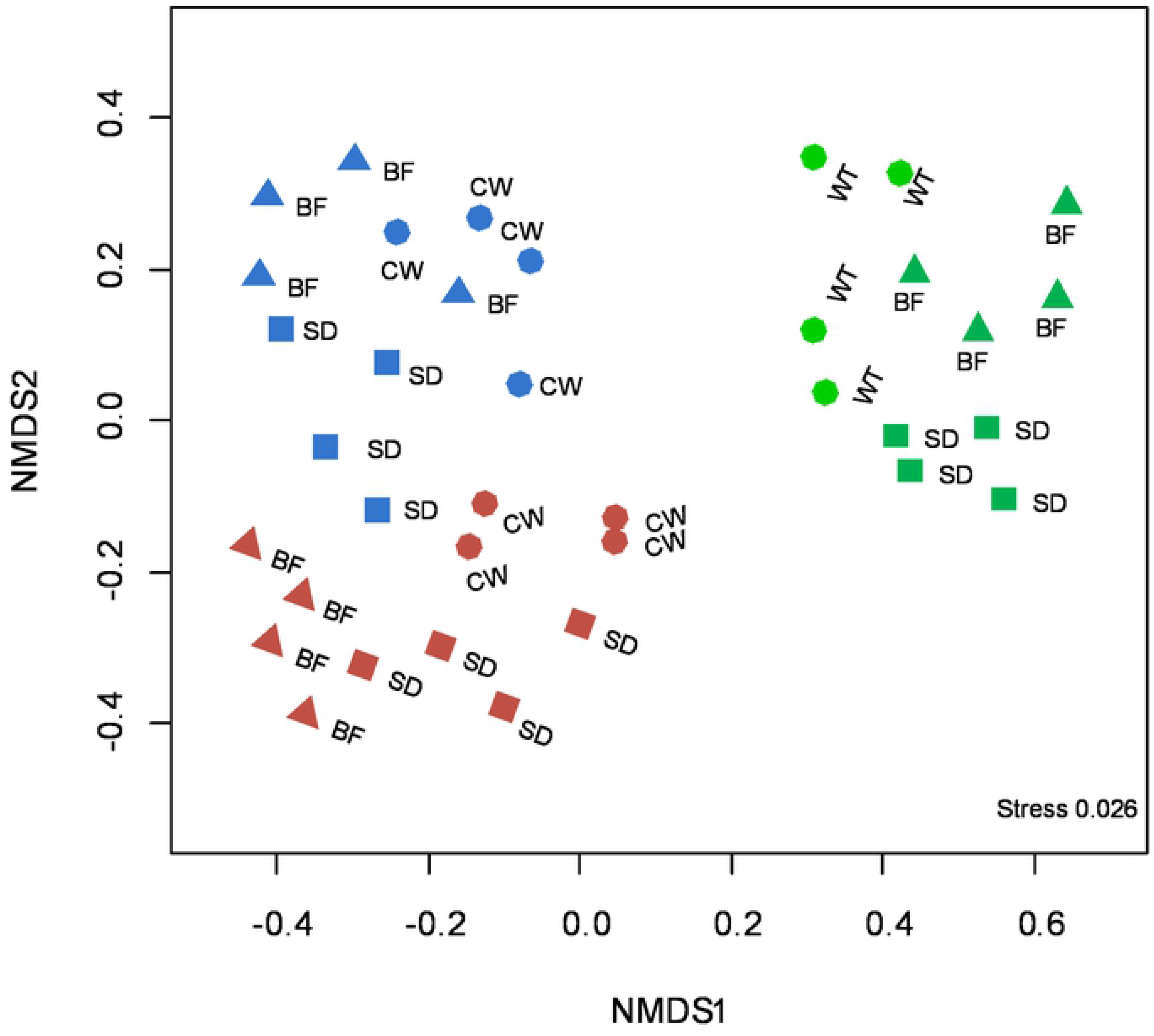
Non-metric multidimensional scaling plot based on the Bray-Curtis dissimilarity showing the relationship between the samples in each treatment. Shapes in triangle, circles, and squares represent biofilm, constructed-wetland, and sedimentation treatments respectively. Colors in brown, blue and green represent samples at the surface, middle and bottom sediment depth in each system respectively

**Table 2.**
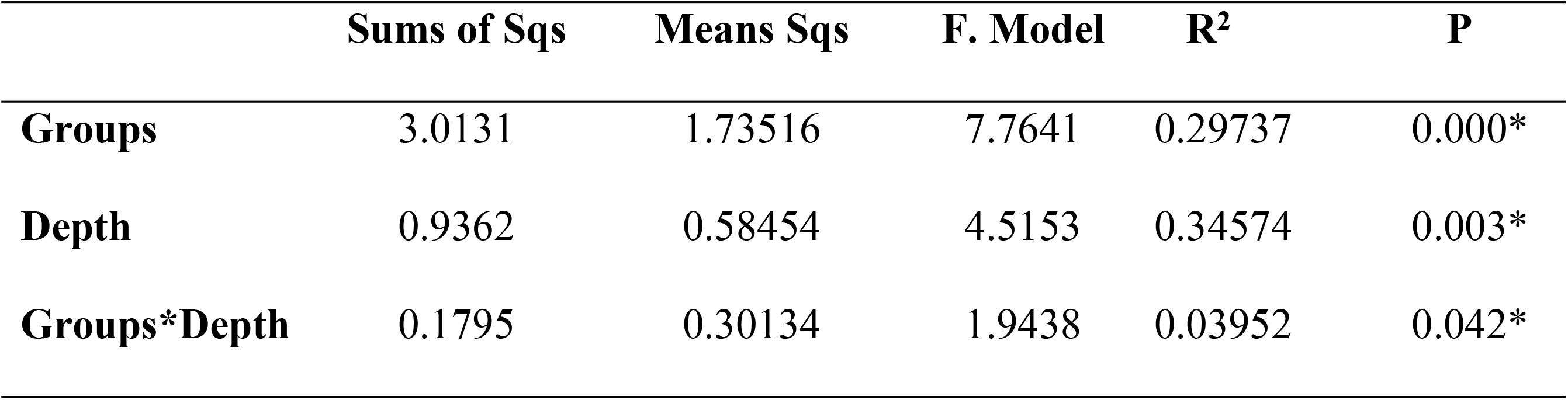
16S rRNA sediment microbial community distributions, structure, and composition determined by a permutational multivariate analysis of variance (PERMANOVA, Adonis function) among the treatments.

|  | Sums of Sqs | Means Sqs | F. Model | R<sup>2</sup> | P |
| --- | --- | --- | --- | --- | --- |
| **Groups** | 3.0131 | 1.73516 | 7.7641 | 0.29737 | 0.000* |
| **Depth** | 0.9362 | 0.58454 | 4.5153 | 0.34574 | 0.003* |
| **Groups\*Depth** | 0.1795 | 0.30134 | 1.9438 | 0.03952 | 0.042* |

### Microbial community and environmental variables

To explore the relationship between the sediment-microbial community and environmental factors, a correlation analysis was performed based on CCA. The analysis showed significant correlations between microbial community composition (genus level) and the environmental factors (Mantel test, p<0.05). The CCA showed the two components of the graph jointly explained 78.89% (axis 1: 41.37% and axis 2: 29.52%) of the total sediment microbial community variance, implying that physicochemical factors and bacterial community composition/structure had a substantive influence over the other. Generally, nine environmental variables were significantly associated with the bacterial community among the treatments (Fig 5). The weakest correlation was observed between SRP and NO_2_^−^-N and the communities (sedimentation). A significant correlation between *Desulphobacterales, Nitrospira*, and *Clostridia* taxa and the variables TN, TOC, and TP were evident. Significant correlations between *Nitrospira* and TOC, TN, and SRP (biofilm) and TAN in the constructed-wetland samples were observed. Furthermore, significant correlations between *Desulfomicrobium, Cytophagales*, and *Planctomyces* and NO_2_^−^-N, and TP in the sedimentation samples were found. Whilst *Ferimonas* and *Verrucomicrobium* were positively correlated with TOC, *Burkholderiales* positively correlated with TAN, TP, and SRP especially in the constructed-wetland samples (Fig 5).

**Fig 5.**
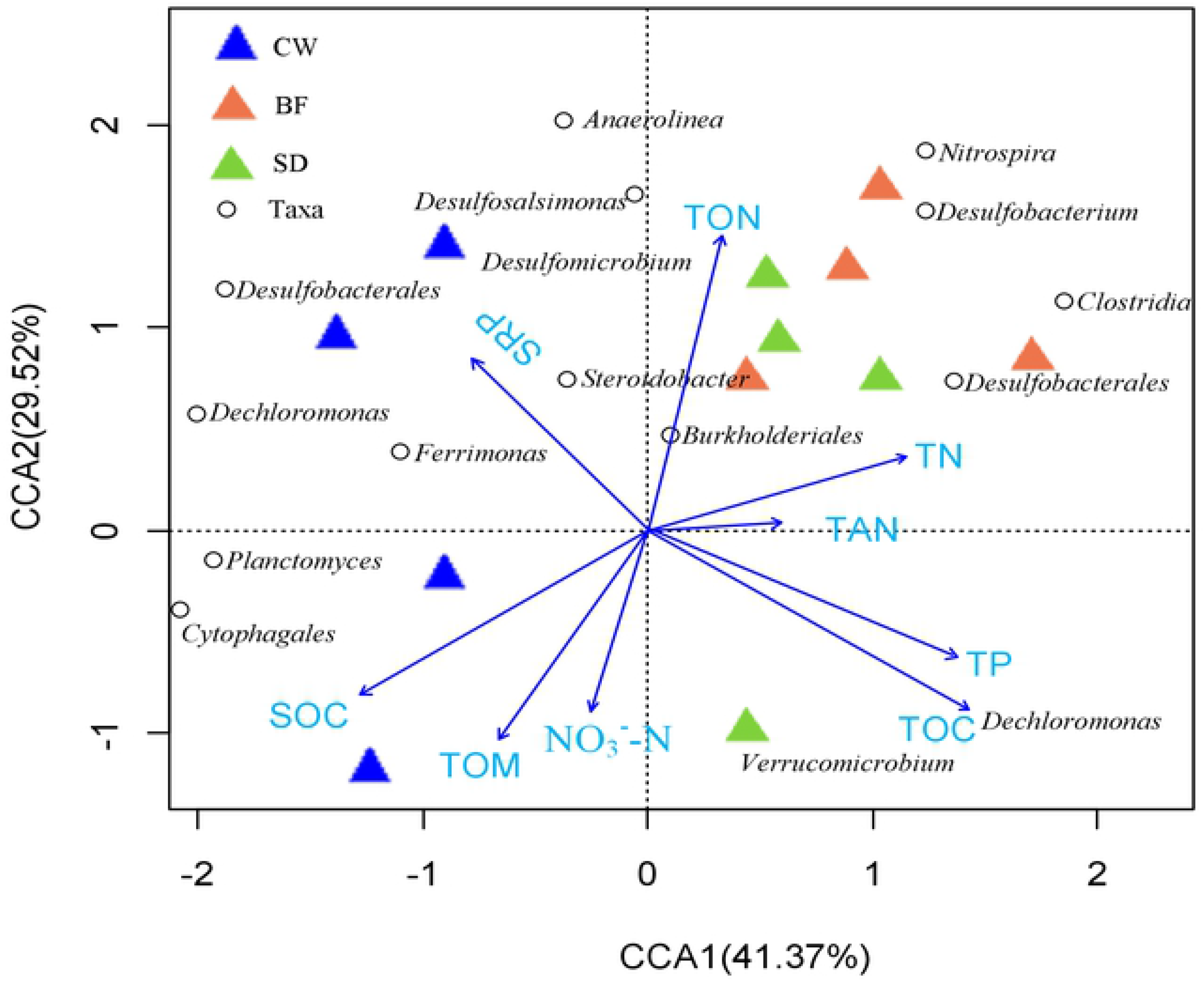
Canonical Correspondence Analysis ordination plot showing the relationships between bacterial community and physicochemical variables in the three treatments of the aquaculture tail-water treatment system. Samples collected from the surface, middle, and bottom of each system are designated with blue, orange, and green triangles, respectively. The abbreviations indicate sediment oxygen consumption (SOC), total-organic matter (TOM), organic nitrogen (TON), ammonia nitrogen (TAN), phosphate (TP), nitrate (NO_3_^−^-N), nitrite (NO_2_^−^-N) and sediment reactive phosphate (SRP)

### Sediment microbial functional groups distributions

Using FAPROTAX the analysis revealed a comparative number of various specific metabolic functional groups/pathways (e.g., chemo-heterotrophy, nitrate-ammonification, nitrification, denitrification, Fig 6) associated with the 16S rRNA genes. Different functional groups involved in nitrogen transformation pathways especially in the surface (0-2 cm) and middle (2-4 cm) cores were predicted suggesting elevated nitrogen mineralization activities. The relative abundance of genes mediating denitrification and dissimilatory reduction of nitrate to ammonia were mostly higher in the bottom layers (4-8 cm) in all systems. Chemoheterotrophy (29.73±0.11%) was the main metabolic functional group, followed by denitrification (23.51± 0.03%), and complete nitrification (15.05±0.81%). Other promoted functional groups/pathways included aerobic-chemoheterotrophy, sulfate_respiration, nitrate_ammonification, and nitrite_ammonification, especially in biofilm cores. Of all functional groups, groups related to nitrogen-cycling were highly predicted in the constructed-wetland relative to biofilm and sedimentation samples.

**Fig 6.**
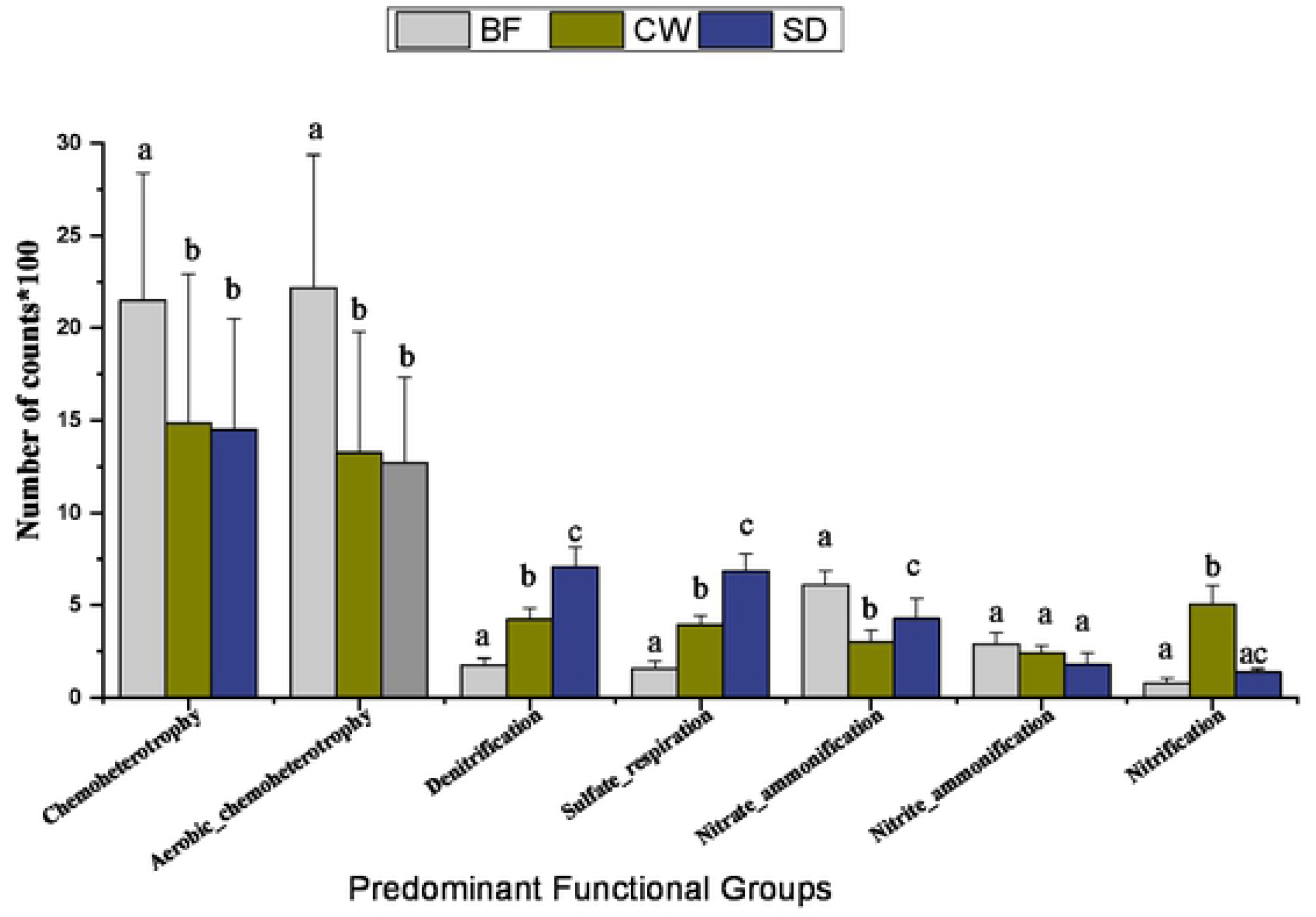
Bar plot of the relative abundance distributions of the predicted predominant functional groups among the treatments as annotated by the FAPROTAX database.

## Discussion

Biofilm, constructed wetland, and sedimentation are potential treatments for improving wastewater and sediment quality [9, 8]. This treatment can improve the ecosystem processes including sediment microbial community activities, distribution, structure, abundance, and successions. The availability and distribution of bacterial communities in the sediment can help to improve and optimize the bioremediation process [8, 41, 42]. This study indicates that a couple of ecological activities including microorganism distribution patterns, nutrient dynamics, and contents of the OM were significantly influenced by the treatments

### Microbial community composition and distribution

The sedimentary bacterial community compositions and structure among the treatments were differently distributed (Figs 3, 4; Table 2). Biofilm and constructed wetland treatments supported more abundant and distinct microbial communities especially at the surface layers (Fig 2). Particularly, phyla such as Proteobacteria and Acidobacteria were most abundantly distributed across all samples, with the highest relative abundance been recorded in biofilm samples relative to constructed-wetland and sedimentation suggesting an elevated mineralization hence nutrient fluxes (Fig 1). This suggests that the treatments probably created varying ecological conditions that accelerate microbial activities and multiplication. The *Proteobacteria, Bacteroidetes, Acidobacteria, Deltaproteobacteria*, *Clostridia, Firmicutes Gammaproteobacteria*, and *Bacilli* were among the most dominant phyla with approximately 37% and 43% bacterial composition in constructed-wetland and biofilm respectively compared sedimentation (19%) (Fig 4). Some studies suggest that soil microbial distribution can be regulated by the different vegetation types [43, 44]. In this study, microbial diversity was highly distributed especially within the constructed-wetland subsystem (Figs. 2A-C, 3). Bodelier, [45] and Lukwambe *et al*. [9] wetland rhizospheres are oxic-habitats created by the roots’ aeration and can markedly affect the diversity of the wetland’s heterotrophic biota and activate nutrient fluxes.

There is a substantial interaction among the constructed wetland plants, microorganisms, and contaminants supported by their complex rhizosphere system [46]. Besides, the plants roots forming the constructed-wetland harbor/store useful nitrifying-denitrifying bacteria [47]. This is evident especially with the rhizosphere of emergent aquatic plants, where the plant roots provide a favorable habitat and exudate the growth of various microbes responsible for sediment reworking including sediment nitrogen content transformation [48, 49]. A profound number of *Proteobacteria, Chloroflexi*, and *Acidobacteria*, were observed within the biofilm and constructed wetland, this may suggest the presence of elevated mineralization activities which affect the succession and stability of the bacterial community [50, 51, 52]. Also, a biofilm environment can potentially improve bacterial communities distributions [53]. Based on this observation, we can deduce that both biofilms and constructed wetlands favored more bacterial community related to organic wastes degradation relative to other sedimentation systems.

Its known remediation measures by using constructed-wetland, biofilm, and sedimentation are known to influence the aquatic ecosystem biodiversity due to improved sediment conditions [29]. The sedimentary ecological niches created by each system differently affect the biotic and abiotic characteristics, thereby resulting in the change of the aquatic microbial diversity and functional diversity [54].

### Biogeochemical fluxes and functional microbial community

In the current study, constructed-wetland showed higher SRP, NO_2_^−^-N, NO_3_^−^-N, and TAN flux rates relative to other treatments (Fig 1). This release pattern is ascribed to promoted physicochemical-microbial mediated activities such as mineralization, nitrification-denitrification, and redox reaction [8, 19]. Literatures show that the root system of the constructed-wetland plants has rhizomes that aerate the sediment potentially resulting in increased dissolved oxygen which promotes microbial assemblages and nitrification-denitrification activities [9, 55, 56]. In this study, we found several bacterial taxa associated with ammonium oxidizing bacterial (AOB, e.g., *Nitrospira*) and nitrite-oxidizing bacteria (NOB, *Nitrospina, Nitrosomonas*) and sulfate-reducing bacteria (SRB, *Desulfatibacillum* and *Desulfobacterium*) which contribute to effluent degradation and material transformation [**26**, 56]. These species were enriched in both constructed wetland and biofilm indicating that the elevated nutrient fluxes were probably due to enhanced bacterial activities such as organic matter mineralization. On the other hand, putatively performing dissimilatory nitrate reduction to ammonia taxa were about 2.5- to 3-fold more in biofilm and constructed-wetland suggesting an increased mineralization activity including nitrification. Normally nitrification process is facilitated by both AOB and NOB bacterial [57]. Under the presence of oxygen microbial nitrogen transformation is supported. Vila-Costa *et al*. [58] reported that macrophyte species with high root oxygen release capacity may enhance the diversity and activity of ammonia oxidizers leading to increased nitrogen content transformation.

Lower TAN fluxes were observed in the biofilm treatment system suggesting increased ammonia utilization by nitrifying bacteria such as *Nitrosomonas* and *Nitrobacter*. These group of bacterial are reported to reduce excess nitrogenous content in the sediment [19]. In our study, several bacterial functional groups related to biogeochemical nutrient metabolism, cycling, and degradation were discovered (Fig 6). The expression of chemoheterotrophy, aerobic-chemoheterotrophy, and denitrification microbial functional groups was significantly higher in the constructed wetland than biofilm and sedimentation. This implied that constructed-wetland best enhanced the activities related to effluent degradation that led to increased nutrient transformation and fluxes. This as well suggests that the genes associated with different biogeochemical functions were favored and enhanced. Sediment nitrogen fixation, nitrification, denitrification, ammonification, and other major nitrogen transformation processes are mediated by soil bacteria [56, 59]. The sedimentary nitrogen cycle can be improved by biological nitrification and denitrification pathways leading to healthy environmental ecosystems. For example, we observed an increase in the absolute content of functional group related *Acinetobacter* (*Moraxellaceae*), which is responsible for detoxification of different pollutants, such as degradation of aromatic compounds [60]. The increased SRP flux rate from the sediment into the water (biofilm, Fig. 1B) indicates organic matter transformation could have been promoted by the bacterial community. Ki *et al*. [56] indicated that organic wastes can be decomposed into soluble reactive phosphate by the SRB bacteria, such as *Desulfobacterium, Desulfatibacillum, Desulfomicrobium*, and *Desulfosalsimonas*, which were most evident in both biofilm and constructed-wetland.

### Bacterial community and sediment organic contents

The content of TON, TOC, TP, and TOM varied significantly among the treatments (Table 1). The observed distribution trend was likely due to improved bacterial community activities associated with mineralization, such as nitrification-denitrification. Sediment nitrogen fixation, nitrification, denitrification, ammonification, and other major nitrogen transformation processes are mainly mediated by soil bacteria [59]. The expression of *Dechloromonas, Steroidobacter*, and *Novosphingobium* among the treatments are likely to support the denitrification process and strengthen the physicochemical-microbial interactions. For instance, *Dechloromonas* has been described as denitrifers that produce nitrogen gas as a reduced nitrogen product [61]. Fabian *et al*.[62] stated that denitrification can also be fueled by the presence of *Steroidobacter* in the sediments. Generally, the lower TN and OM in biofilm and constructed wetland over the sedimentation is probably due to the TAN transformation through microbial oxidation to NO_3_^−^-N and NO_3_^−^-N. The majority of nitrogen content reduction in wetlands is believed to result from the microbial coupled nitrification-denitrification interactions and uptake by the wetland plants [63]. The 16S rRNA sequencing result showed that taxa such as *Firmicutes* and *Nitrospinae* were differentially enriched among the treatments a phenomenon that may have attributed to the reduced level of the organic contents.

Additionally, CCA indicated strong relationships between the bacterial communities and physicochemical factors (Fig 5). Among the treatments, nine environmental variables (TAN, NO_2_^−^-N, NO_3_^−^-N, SRP, TN, TP, TOC, OM and, SOC) correlated more closely with microbial community groups. The correlations between bacterial communities and nutrient fluxes and organics were moderately high (ordination axis 1 = 58.1%, axis 2 = 42.7% of the total variation). The constructed-wetland and biofilm had more affiliated taxa linked with physicochemical variables relative to sedimentation indicating a greater association between the functional genera among the two treatments. This result is similar to Wu *et al*. [64], Lukwambe *et al*. [29] and Ki *et al*. [56] who reported a substantial correlation among the bacteria and nitrogen transformation. In similar patterns, CCA results revealed that TAN, TP, TOC, and TN contents were factors that strongly correlated with *Desulfomicrobium* (surface sediment) and *Chloroflex* (deeper sediment, biofilm) while SOC, TP, and SRP mostly correlated with *Ferrimonas, Burkholderia, Dechloromonas*, and *Desulfomicrobium*, especially in constructed-wetland. This can be supported by a previous study [56, 65] which indicated *Desulfomicrobiuim* and *Methylobacter* had strong association with TAN resulting in reduced organic contents.

### Conclusions

This study investigates the distributions of the bacterial community, nutrient-fluxes, and organic matter contents in a comprehensive aquaculture tail-water treatment system. The study showed that the treatments differently improved the sediment bacterial dynamics (community structure, diversity, and composition), elevated nutrient dynamics and fluxes, and reduced organic matter contents. Microbial groups associated with AOB, NOB, SRB were enriched in the constructed wetland and biofilm but so within the 0-4 cm sediment depth. The chemoheterotrophy, aerobic-chemoheterotrophy, denitrification, and nitrification were the most dominant functional groups of all treatments but especially in the constructed wetland. The TAN, NO_2_^−^-N, and NO_3_^−^-N nutrient flux rates across the sediment-water interface were higher in constructed-wetland than in biofilm and sedimentation subsystems. The constructed-wetland and biofilm had lower organic effluents and better sediment conditions relative to sedimentation. Our study suggests that bacterial diversity and structure were highly improved especially under constructed-wetland. Whereas biofilm best promoted the bacterial community composition relative to other treatments.

## Acknowledgments

This study was supported by National Key R & D Program of China (2020YFD0900201), the Zhejiang Public Welfare Technology Research Program of China (ZPWTP) (LGN18C190008) and the K.C. Wong Magna Fund in Ningbo University

